# SemiBin2: self-supervised contrastive learning leads to better MAGs for short- and long-read sequencing

**DOI:** 10.1101/2023.01.09.523201

**Authors:** Shaojun Pan, Xing-Ming Zhao, Luis Pedro Coelho

## Abstract

**Motivation:** Metagenomic binning methods to reconstruct metagenome-assembled genomes (MAGs) from environmental samples have been widely used in large-scale metagenomic studies. The recently proposed semi-supervised binning method, SemiBin, achieved state-of-the-art binning results in several environments. However, this required annotating contigs, a computationally costly and potentially biased process.

**Results:** We propose SemiBin2, which uses self-supervised learning to learn feature embeddings from the contigs. In simulated and real datasets, we show that self-supervised learning achieves better results than the semi-supervised learning used in SemiBin1 and that SemiBin2 outperforms other state-of-the-art binners. Compared to SemiBin1, SemiBin2 can reconstruct 8.3%–21.5% more high-quality bins and requires only 25% of the running time and 11% of peak memory usage in real short-read sequencing samples. To extend SemiBin2 to long-read data, we also propose ensemble-based DBSCAN clustering algorithm, resulting in 13.1%–26.3% more high-quality genomes than the second best binner for long-read data.

**Availability and Implementation:** SemiBin2 is available as open source software at https://github.com/BigDataBiology/SemiBin/ and the analysis scripts used in the study can be found at https://github.com/BigDataBiology/SemiBin2_benchmark.

## 1 Introduction

Microorganisms live in all environments on Earth and play essential roles in human health, agriculture, food, climate change, and other processes^1^. Considering the difficulty of culturing microorganisms, metagenome sequencing is widely used to study microorganisms^2^. Due to the low cost of short-read sequencing, large compendia of metagenome-assembled genomes (MAGs) have been built, expanding the known diversity of bacteria in human-, animal-associated, and environmental habitats^3–7^. Despite the success of short-read sequencing, it often fails to reconstruct repeated elements^8,9^. Recently, long-read sequencing technologies, such as PacBio and Oxford Nanopore which address this limitation, have started to become popular^10–12^.

Whether using short- or long-read sequencing, assembling large contiguous sequences (contigs) from individual reads is the first step in recovering MAGs. Metagenomic binning is a clustering problem, namely grouping together contigs inferred to originate from the same organism to reconstruct MAGs^13^. Several binning methods have been proposed. Most existing unsupervised metagenomic binners, such as Canopy^14^, SolidBin^15^, MetaBAT2^16^, MaxBin2^17^, MetaDecoder^18^ reconstruct bins using *k*-mer frequencies and abundance information. Recently, deep learning has been applied to this problem. VAMB^19^ uses deep variational autoencoders to encode *k*-mer and abundance features prior to clustering. SemiBin (henceforth, SemiBin1)^20^ implements a semi-supervised approach to learn an embedding by contrastive learning with information from reference genomes, and it achieved state-of-the-art binning results across several habitats, and different binning modes^20^.

Nevertheless, semi-supervised learning has two drawbacks: (1) SemiBin1 requires using a contig annotation tool such as MMseqs2^21,22^ to generate cannot-link constraints, which significantly increases the running time and peak memory usage of the binning process^20^; (2) limitations of the reference genome databases lead to bias (some genomes cannot be annotated and will not be covered in the cannot-link constraints)^20^ (see Supplementary Table 2). Binning can be performed per-sample or using multiple samples at once, a setting termed multi-sample^19^. For single-sample binning, embedding models can be be pretrained from large collections of samples, which can alleviate the annotation bias problem and, given the fact that models can be reused, the computational costs are amortized. On the other hand, for multi-sample binning, models need to be relearned for each binning task (as they depend on the number of samples used). So here we focus on improving multi-sample binning, which was shown to lead to the highest number of recovered high-quality MAGs^19,20^. In particular, we propose a self-supervised binning method that does not require reference genome annotation.

Another limitation of existing binners is that most of them are not optimized for long-read sequencing data. Even though they can be used, assemblies from long-read sequencing are significantly different from short-read sequencing (see Supplementary Table 1) and results will be sub-optimal. Thus, SemiBin2 proposes an ensemble-based DBSCAN clustering algorithm to extend to long-read data. We compared it to the other binners proposed for long-read sequencing data, such as LRBinner^23,24^ and the recently-proposed GraphMB^25^, and show that it outperforms them.

We have shown that SemiBin2 obtains state-of-the-art binning results on the short- and long-read sequencing data and needs much less computational resources (less running time and peak memory usage) than SemiBin1.

## 2 Materials and methods

### 2.1 The SemiBin2 algorithm

We developed SemiBin2, a self-supervised contrastive deep learning-based contig-level binning tool for short- and long-read metagenomic data (see Fig. 1). As in SemiBin1, SemiBin2 uses must-link and cannot-link constraints to learn a feature embedding prior to clustering^20^. However, SemiBin2 improves on SemiBin1 by using self-supervised learning to obtain cannot-link constraints: randomly sampled pairs of contigs are assumed to contain a cannot-link between them.

**Fig 1.**
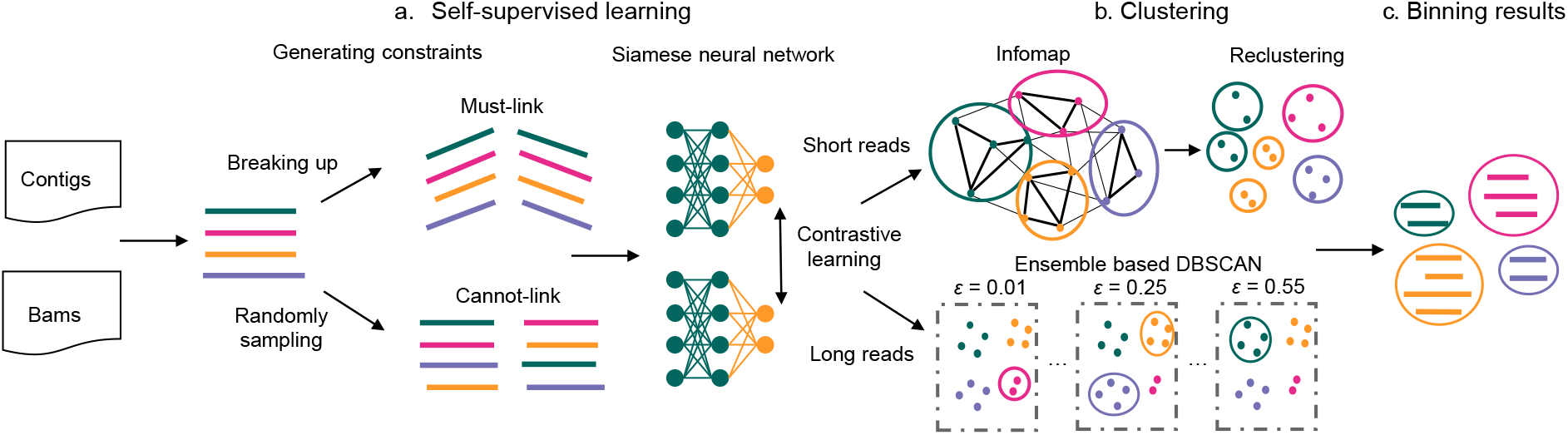
Overview of the SemiBin2 pipeline. **a**, Self-supervised learning, including two steps: constraint generation and the siamese neural network. Generating must-link constraints is done by breaking up longer contigs and cannot-link constraints by random sampling. Then, a deep siamese neural network is used to learn a better embedding from the inputs. **b**, For short-reads, the Infomap algorithm is used to obtain preliminary bins from the sparse graph generated from the embeddings, followed by weighted *k*-means to recluster bins whose the mean number of single-copy genes is greater than one. For long-reads, SemiBin2 runs DBSCAN with different values of the *ε* parameter with embeddings as inputs and integrates the results based on single-copy genes. **c**, Output the final binning results larger than a user-definable threshold (default 200kbp).

The clustering approach used depends on the type of data. For short-read data, the same community detection-based method employed in SemiBin1 is used^26^. For long-read data, however, a novel ensemble-based DBSCAN method is used (see below).

#### 2.1.1 Preprocessing

Every contig is represented by its tetramer frequencies and its estimated abundance (the average number of reads per base mapping to the contig). Depending on the numbers of samples and sequencing technology, SemiBin2 uses different ways to process the abundance values. Assuming the original abundance value is *a* and the number of samples is *N*, SemiBin2 processes the abundance as follows:

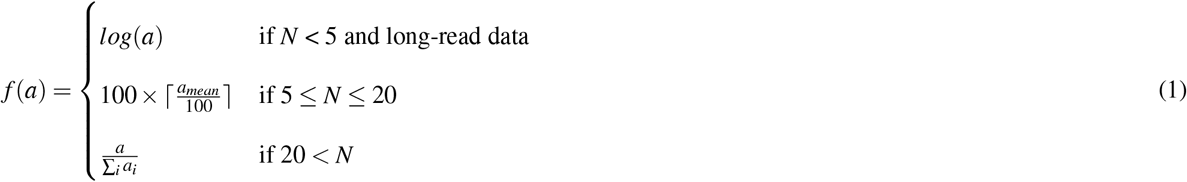

*a*_*mean*_ is the average of all abundance values. The inputs to the deep learning model are the *k*-mer frequencies and preprocessed abundance values.

#### 2.1.2 Self-supervised contrastive learning

SemiBin2 uses the same approach as SemiBin1 to generate must-link constraints, namely simulating the break up of longer contigs. For the generation of cannot-link constraints, SemiBin1 used taxonomic annotations which carried large computational costs (including high memory requirements which limited the accessibility of the method). SemiBin2 uses a self-supervised approach, randomly sampling pairs of contigs and treating them as cannot-link pairs.

To control the training time and the ratio between must-link and cannot-link constraints, SemiBin2 limits the number of cannot-link constraints used in training to min(*m*, 4000000), where *m* is the number of must-link constraints.

Then, SemiBin2 uses a deep siamese neural network^27^ to implement a contrastive learning method and learns a better embedding from the must-link and cannot-link constraints. The first two layers of the network are followed by a batch normalization^28^ layer, a leaky rectified linear unit^29^ layer and a dropout layer^30^:

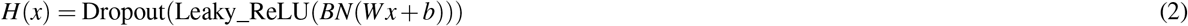

where *H*(*x*) is the output of the neural network, *x* is the inputs, *W, b* is the parameter of the neural network.

SemiBin2 uses a supervised contrastive loss to classify the must-link (positive label) and cannot-link (negative label) constraints. This is unlike SemiBin1, which combines this loss with an unsupervised reconstruction loss. In particular, the conterastive loss function used is:

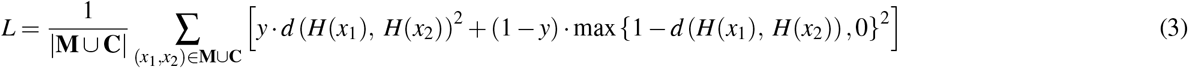

where *x*_1_, *x*_2_ are input features, **M** denotes all must-link constraints, **C** denotes all cannot-link constraints, *d* the Euclidean distance, and y is the label such that 1 denotes *x*_1_ and *x*_2_ contain a must-link constraint between them while 0 denotes they contain a cannot-link constraint. The loss function is minimized with Adam optimization algorithm^31^.

#### 2.1.3 Clustering of short- and long-read data

After obtaining the contig embeddings from the deep learning model, a clustering algorithm is used to obtain the final bins. For short-read data, SemiBin2 uses the same clustering method used in SemiBin1^20^: a community detection-based clustering method to generate preliminary bins and a weighted *k*-means algorithm to find the final binning results by reclustering, informed by single copy genes^32^.

However, preliminary testing showed that this is not suitable for long-read data as assemblies from long-read data have different properties compared to assemblies from short-read data. In particular, long-read contigs are fewer and much longer (see Supplementary Table 1). This results in some genomes consisting of a small number of contigs (even a single contig). The existing tools are mostly optimized for short-read data and do not work very well when applied to long-read data.

Therefore, for long-read data, SemiBin2 employs a ensemble-based DBSCAN^33^ algorithm to bin contigs. DBSCAN is a density-based method where a user-tunable parameter (*ε*) can be used to influence the size of resulting clusters. A smaller *ε* value will lead to smaller clusters and a larger *ε* value will lead to larger ones. SemiBin2 uses the implementation of DBSCAN in scikit-learn^34^ and runs DBSCAN with *ε* value equals to 0.01, 0.05, 0.1, 0.15, 0.2, 0.25, 0.3, 0.35, 0.4, 0.45, 0.5, and 0.55. Then SemiBin2 integrates the results of these runs based on the single-copy genes that have been used in other tools^18,25^. In particular, SemiBin2 uses 107 single-copy genes^32^ to estimate the completeness, contamination, and F1-score of every bin.

Assume we find *G* genes and *N* nonredundant genes in the bin.

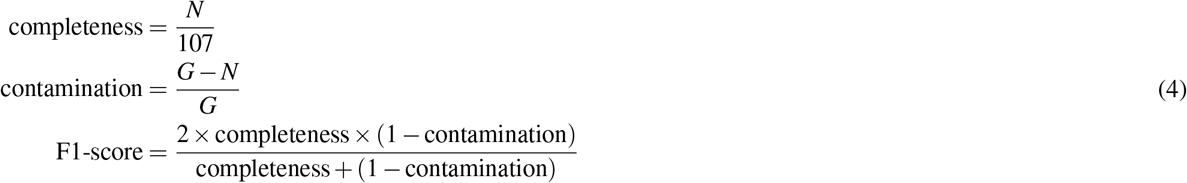

Based on these metrics, SemiBin2 uses a greedy algorithm to select the final bins: at each step, the best bin is selected and all its contigs are removed from further consideration until no more bins can be found that fulfill the minimal quality criteria (see Algorithm 1).

##### Algorithm 1 Integrate all runs of DBSCAN

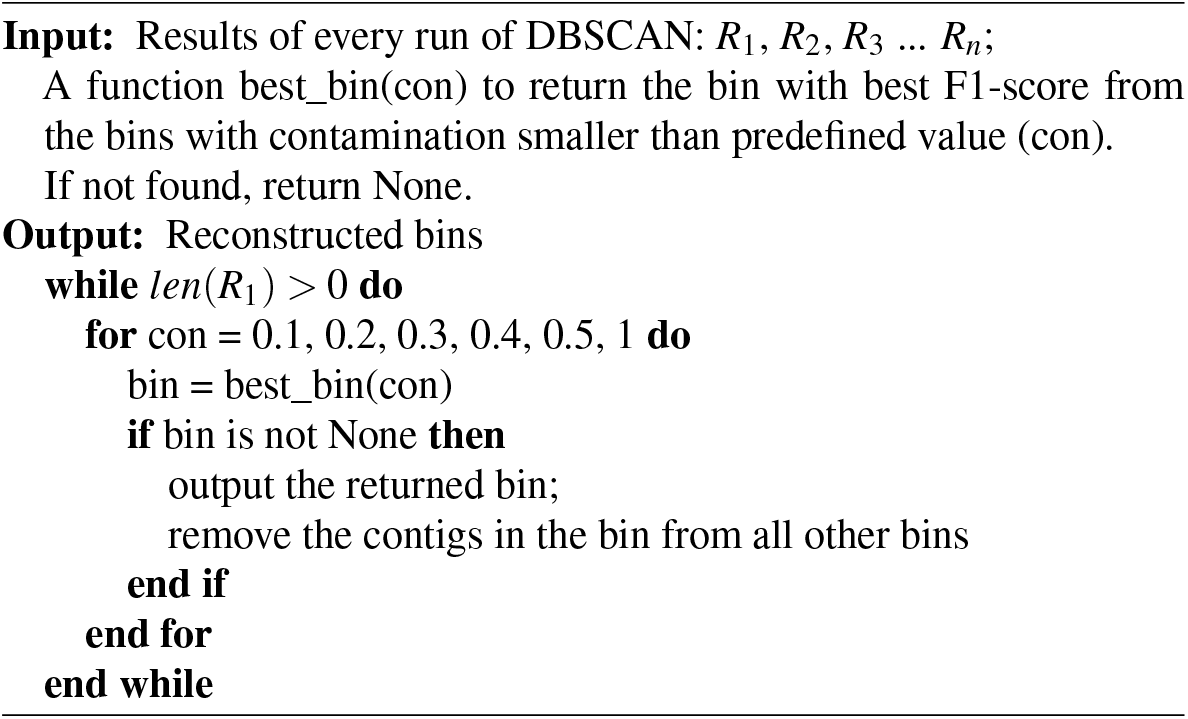

### 2.2 Data used for benchmarking

For benchmarking, we used 5 simulated datasets from the CAMI II challenges, real data from 4 short-read sequencing projects and from 3 long-read sequencing projects. CAMI II datasets comprise 5 human-associated environments: airways (10 samples), gastrointestinal (10 samples), oral (10 samples), skin (10 samples) and urogenital (9 samples). Every environment has short-read simulated datasets and PacBio (long-read) simulated datasets.

We used the same 4 short-read sequencing projects and the same assemblies previously used to evaluate SemiBin1: human gut (82 samples)^35^, dog gut (129 samples)^36^, ocean (109 samples)^37^, and soil (101 samples)^38^. For long-read datasets, we used three projects: PRJCA007414 (human gut, 3 PacBio-HiFi samples, 3 Nanopore R9.4 samples), PRJNA595610 (sheep gut, 2 PacBio-HiFi samples)^10^ and PRJEB48021 (activated sludge from an anaerobic digester, 1 PacBio-HiFi sample, 2 NanoPore R9.4.1 samples and 1 Nanopore R10.4 sample)^12^ (see Supplementary Table 3). Long-reads were assembled with Flye^39^ (version 2.9-b1768, *–pacbio-hifi* for assembling PacBio-HiFi reads, *–nano-raw* for assembling Nanopore reads from PRJCA007414 and *–nano-hq* for assembling Nanopore reads from PRJEB48021). For correction of the assemblies from Nanopore reads, we used the tools used in the original studies. For the correction of assemblies from PRJCA007414, we used Pilon^40^ (version 1.24) and for assemblies from Nanopore R9.4.1 from PRJEB48021, we runned Medaka (version 1.7.1, *-m r941_min_sup_g507*) and Racon^41^ (version 1.5.0, used two times, one round using long-reads and one round using short-reads). For short-reads, we used Bowtie2^42^ to generate the mapping (BAM) files and for long reads, we used Minimap2^43^ (version 2.24-r1122, *-x map-hifi* for PacBio-HiFi reads and *-x map-ont* for Nanopore reads).

The simulated CAMI II datasets can be downloaded from https://data.cami-challenge.org/participate. The short-read datasets used in the study can be found in SemiBin1^20,35–38^. The long-read datasets used are publicly available in the NGDC with the study accession PRJCA007414 and in the ENA with the study accessions PRJNA595610 and PRJEB48021.

### 2.3 Methods used in benchmarking

For short-read datasets with multi-sample binning, we compared to VAMB (version 3.0.7, *-m 2000*)^19^ and SemiBin1 (version 1.0.0)^20^, existing binners supporting for multi-sample binning. For long-read datasets, we compared to MetaBAT2 (version 2.15, *–percentIdentity 50*)^16^, VAMB (version 3.0.7, *-m 2000*)^19^, SemiBin1 (version 1.0.0, *–environment human_gut* for PR-JCA007414)^20^, GraphMB (version 0.1.4)^25^, MetaDecoder (version 1.0.13)^18^ and LRBinner (version 2.1)^24^.

To benchmark the value of the embedding specifically, we performed an ablation study whereby we removed the self-supervised learning stope and performed clustering based on the original inputs (a setting we called *NoSemi*).

### 2.4 Evaluation metrics

For simulated datasets, we used gold standard assemblies for binning provided by the CAMI II challenge. We used AMBER (version 2.0.2) to evaluate the results of the simulated datasets (completeness and contamination).

For real datasets, as labels are not known, we evaluated the results using CheckM (version 1.1.9, using lineage_wf workflow)^44^ and GUNC (version 1.0.5). For long-read datasets, as some tools perform binning based on a set of single-copy genes which overlaps with the genes used by CheckM for evaluation, this may overestimate the quality of the outputs. Thus, we used CheckM2^45^ (version 0.1.3) which is based on machine learning to estimate the completeness and contamination for long-read datasets.

For simulated datasets, we defined high-quality bins as those with completeness > 90% and contamination < 5%. For short-read datasets, we defined high-quality bins as those with completeness > 90%, contamination < 5%, and passing the chimeric detection of GUNC. For long-read datasets, we termed the high-quality bins defined before as near-complete bins and defined high-quality bins as those with completeness > 90%, contamination < 5%, passing the the chimeric detection of GUNC and having at least one 23S, 16S, 5S rRNA genes and 18 distinct tRNAs. We used Barrnap (version 0.9, https://github.com/tseemann/barrnap) and tRNAscan-SE (version 2.0.9)^46^ to detect these genes.

## 3 Results

### 3.1 Self-supervised learning reduces resource usage and improves results

When comparing the cannot-link constraints generated from taxonomic annotation to random sampling on simulated data (where the ground truth is known), we found that taxonomic annotation leads to more accurate constraints. On the other hand, random sampling could cover more genomes (see Supplementary Table 2). Deep learning can be robust to noise^47^, and more genomes covered can provide more information to the model. Thus, it is an empirical question which approach results in better binning.

Compared to semi-supervised learning (used in SemiBin1), self-supervised learning could achieve similar or better results in most of the simulated datasets (see Fig. 2, Supplementary Fig. 1). As simulated datasets are less complex than real-world datasets^20^ and better represented in databases, cannot-link constraints from contig annotations can have high accuracy and coverage, and SemiBin1 and SemiBin2 return comparable results. However, in complex real data, self-supervised learning results in a large improvement in the number of returned high-quality bins (see Fig. 5).

**Fig 2.**
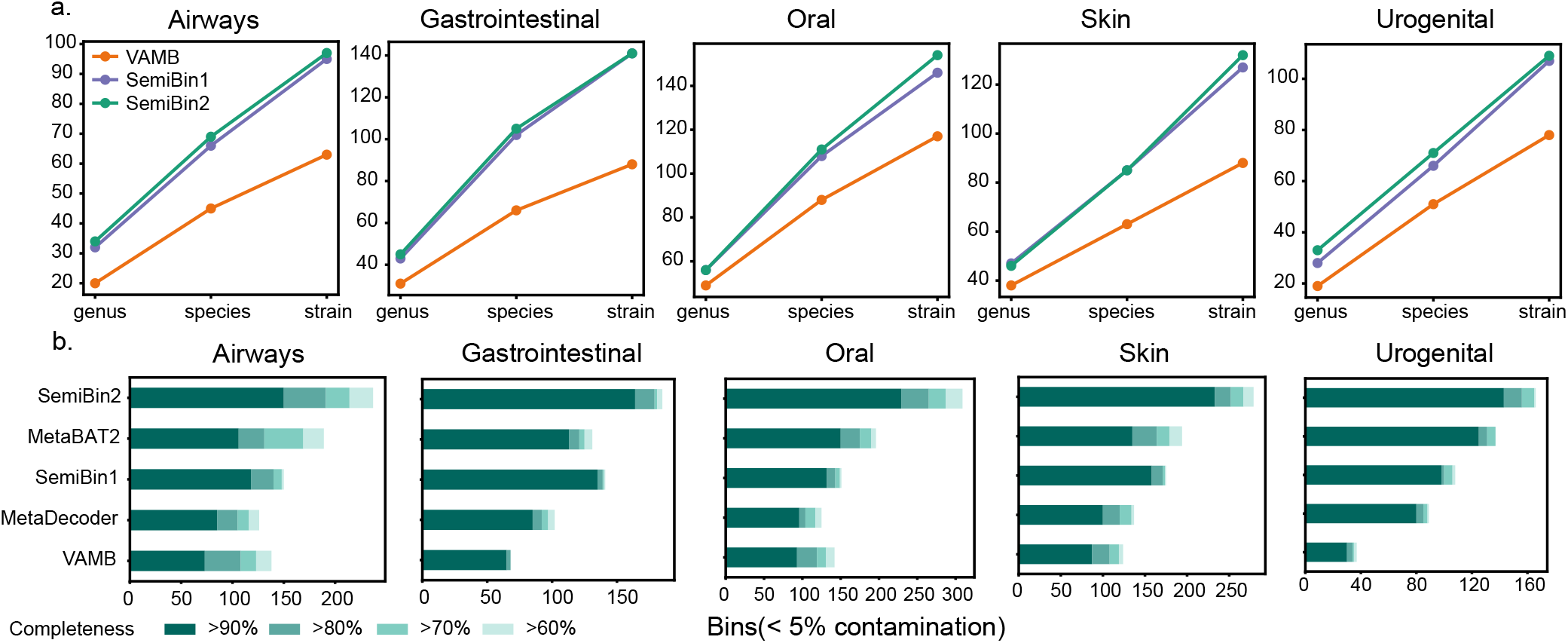
SemiBin2 outperforms other binners with short- and long-read sequencing CAMI II datasets. **a**, SemiBin2 outperformed VAMB and got similar results to SemiBin1 in CAMI II short-read sequencing datasets. Shown is the number of distinct genera, species and strains (see Materials and methods). **b**, SemiBin2 reconstructed many more high-quality bins than other binners in CAMI II long-read sequencing datasets. Shown are the numbers of reconstructed genomes with different completeness and contamination <5%.

In the SemiBin1 workflow, contig annotation, which uses MMseqs2, is the most time-consuming and memory intensive step. Self-supervised learning avoids this step and makes SemiBin2 more efficient than SemiBin1 (see Supplementary Table 5).

To further demonstrate the value of self-supervised constrastive learning, we also compared SemiBin2 to the same pipeline without the deep learning step (termed NoSemi). In the five environments of CAMI II short-read sequencing datasets, SemiBin2 could reconstruct average 21.6% more distinct high-quality strains, 18.4% more distinct high-quality species, and 17.1% more distinct high-quality genera compared to NoSemi (see Fig. 3). For long-read sequencing datasets, SemiBin2 could reconstruct average 31.5% more high-quality bins, showing the effectiveness of self-supervised learning.

**Fig 3.**
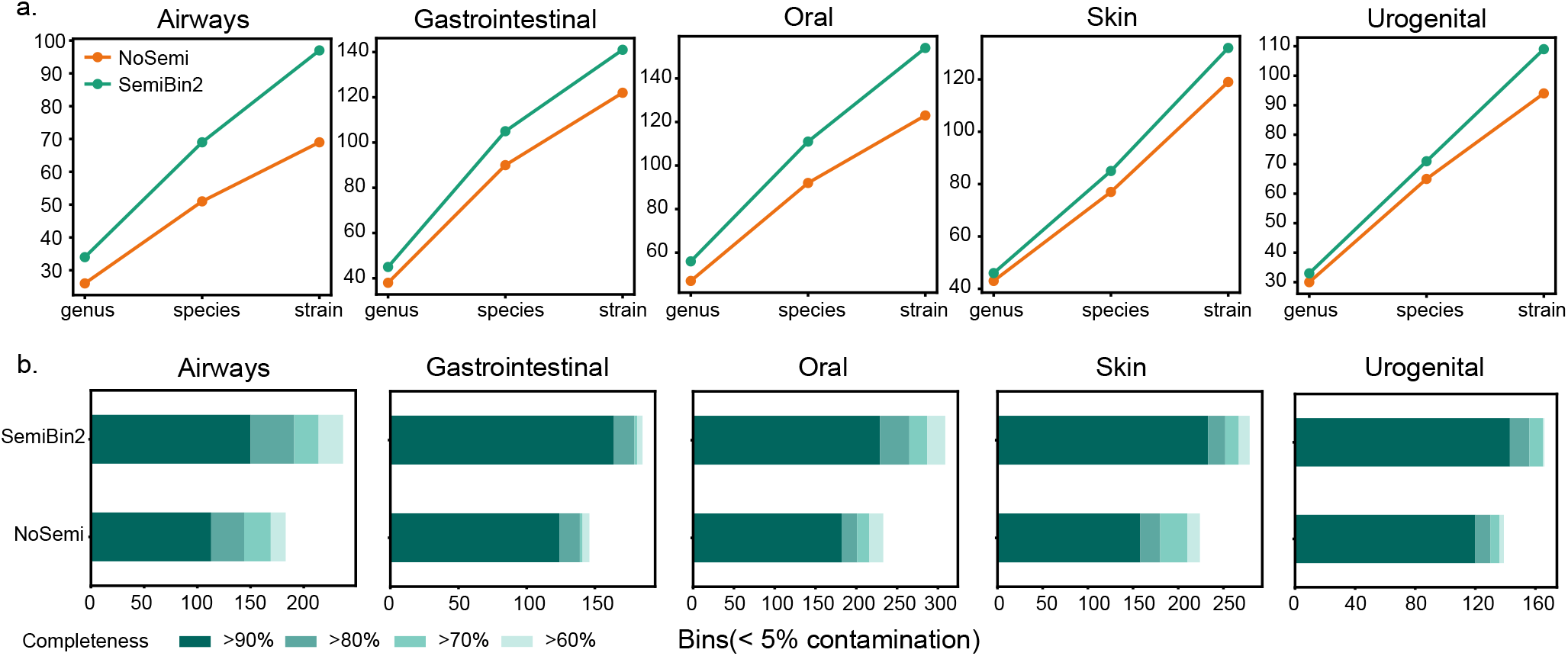
Self-supervised learning improveds binning results in CAMI II datasets. **a**, To evaluate the impact of self-supervised learning, we compare the same pipeline with (SemiBin2) or without (NoSemi) the deep learning step on the simulated CAMI II datasets. **a**, SemiBin2 reconstructed more distinct high-quality genera, species and strains in the five environments from CAMI II compared to the NoSemi version. Shown are the number of distinct genera, species and strains (see Materials and methods). **b**, SemiBin2 reconstructed a more high-quality bins than the NoSemi version. Shown are the numbers of reconstructed genomes with different completeness and contamination <5%.

### 3.2 SemiBin2 outperformed other binners in CAMI II simulated datasets

We compared SemiBin2 to widely used and recently proposed binners on the five environments with short- and long-read datasets from CAMI II. For short-read sequencing datasets, because of the low cost of this technology, there are enough samples to allow us to run binning in multi-sample binning mode, which has been shown to reconstruct the most high-quality bins^19,20^. We compared SemiBin2 with SemiBin1^20^ and VAMB^19^, which are the only two existing binners supporting multi-sample binning. For the long-read datasets, we compared to MetaBAT2^16^, MetaDecoder^18^, VAMB^19^, and SemiBin1^20^. We did not include GraphMB^25^ and LRBinner^24^ in this comparison because we used gold standard assemblies for binning in simulated datasets, and we could not obtain the assembly graph GraphMB requires as input and LRBinner cannot be run with co-assembly binning.

In the short-read datasets, SemiBin2 reconstructed on average 44.8% more distinct high-quality genera (range 14.3–73.7%), 42.5% more distinct high-quality species (range 26.1–59.1%), and 47.1% more distinct high-quality strains (range 31.6– 60.2%) compared to VAMB (see Fig. 2). When compared to SemiBin1, SemiBin2 performed similarly or better, showing the effectiveness of self-supervised contrastive learning and avoiding the time and memory usage required for contig annotations.

For long-read datasets, we proposed an ensemble-based DBSCAN clustering algorithm (see Materials and methods) instead of the community detection approach used for short-read datasets. The ensemble-based DBSCAN clustering algorithm runs DBSCAN with different *ε* values and integrates them using single-copy genes (see Materials and methods). To show that the ensemble step could improve binning results, we compared SemiBin2 (ensemble-based DBSCAN algorithm) to binning with running DBSCAN with a single *ε* value (see Fig. 4). In the airways, gastrointestinal, oral, skin and urogenital environments, SemiBin2 could reconstruct 78.6%, 37.8%, 51.7%, 66.4% and 25.4% more high-quality bins compared to the best result of single DBSCAN running, indicating the ensemble learning could effectively integrate different runs and improve binning results.

**Fig 4.**
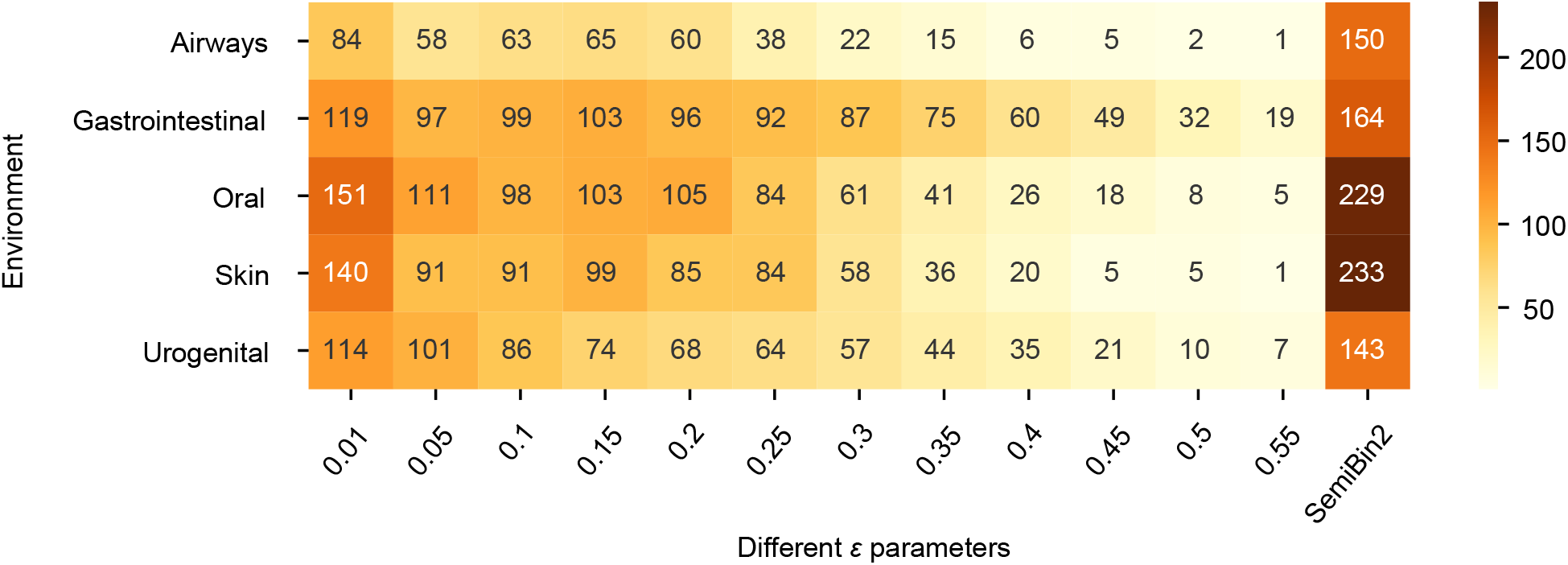
Ensemble-based DBSCAN algorithm improves binning results for long-read data. We compared SemiBin2 (ensemble-based DBSCAN algorithm) to the results of a single DBSCAN run with different *ε* values in CAMI II long-read datasets. Shown is the number of high-quality bins and *SemiBin2* denotes the the result of the ensemble-based DBSCAN algorithm.

Then, we compared SemiBin2 to other binners (see Fig. 2). SemiBin2 outperformed other tools in all environments and reconstructed 27.1%, 21.5%, 52.7%, 47.5% and 14.4% more high-quality bins than the second-best binner in these five environments.

### 3.3 SemiBin2 outperformed other binners in short- and long-read real datasets

We compared SemiBin2 to VAMB^19^ and SemiBin1^20^ with multi-sample binning in short-read sequencing datasets and to MetaBAT2^16^, MetaDecoder^18^, VAMB^19^, SemiBin1^20^, GraphMB^25^ and LRBinner^24^ in long-read sequencing datasets. For short-read sequencing, we used datasets from four environments: human gut^35^, dog gut^36^, ocean^37^ and soil^38^. For long-read sequencing, we chose three long-read sequencing studies: human gut (3 PacBio-HiFi samples, 3 Nanopore R9.4 samples), sheep gut (2 PacBio-HiFi samples)^10^ and activated sludge (1 PacBio-HiFi sample, 2 Nanopore R9.4.1 samples and 1 Nanopore R10.4 sample)^12^. These studies cover different technologies for long-read sequencing that can be used to evaluate SemiBin2 in different situations.

In real data, the true labels of the contigs (*e.g*., which genomes each contig belongs to) are unknown. Thus, we used automated tools (CheckM^44,45^ and GUNC^48^) to evaluate the outputs (see Materials and methods).

SemiBin2 could reconstruct the largest number of high-quality bins for short-read datasets with multi-sample binning. In the four environments, SemiBin2 reconstructed 1678, 3776, 631 and 254 high-quality bins (see Fig. 5). Compared to VAMB, SemiBin2 generated 27.3% (360), 21.5% (669), 44.7% (195) and 229.9% (177) more high-quality bins. Compared to SemiBin1, SemiBin2 reconstructed 8.3% (129), 9.5% (328), 10.7% (61) and 21.5% (45) more high-quality bins. On a sample-by-sample basis, SemiBin2 significantly outperformed VAMB (*P* = 9.7*·*10^−14^ (n = 82), *P* = 4.7*·*10^−22^ (n = 129), *P* = 2.1*·*10^−10^ (n = 109) and *P* = 1.3*·*10^−12^ (n = 101)) and SemiBin1 (*P* = 1.5*·*10^−05^ (n = 82), *P* = 4.1*·*10^−18^ (n = 129), *P* = 8.7*·*10^−05^ (n = 109) and *P* = 5.8*·*10^−04^ (n = 101), *P*-values were computed using Wilcoxon signed rank test, two-sided null hypothesis) (see Fig. 5).

**Fig 5.**
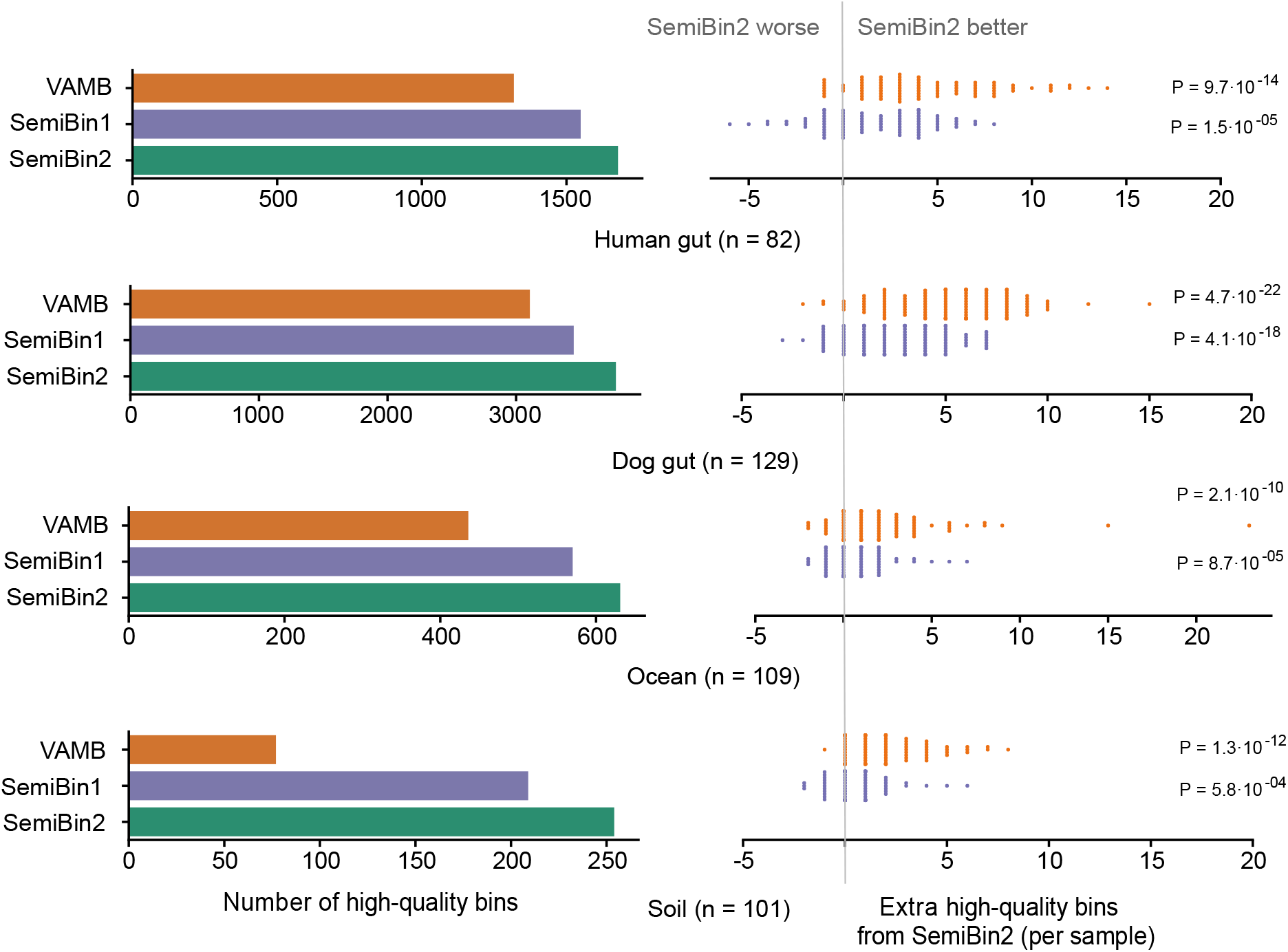
SemiBin2 outperformed other binners in short-read sequencing real datasets with multi-sample binning. SemiBin2 produced more high-quality bins than VAMB and SemiBin1 in the four real datasets with multi-sample binning (lefT) and reconstructed more high-quality bins in most of the samples (right). *P*-values shown are computed using a two-sided Wilcoxon signed-rank test based on the counts of each sample.

When applying them to short-read datasets, the only difference between SemiBin1 and SemiBin2 is the use of semi-supervised vs. self-supervised learning. The results showed that self-supervised learning could learn a better embedding in complex real environments possibly because randomly generated cannot-link constraints cover more genomes (see Supplementary Table 2) and provide more information to the model.

One limitation of SemiBin1 is that it needs to run taxonomic annotation (MMseqs2 by default) to generate the cannot-link constraints. This step requires a lot of computational resources (running time and memory usage). SemiBin2 with self-supervised learning both improves the binning results and addresses this limitation. Taking advantage of self-supervised learning, SemiBin2 needs only *circa* 25% running time on a GPU (the gain when using a CPU is smaller, but still close to 50%; see Supplementary Table 4). More importantly, using either a CPU or a GPU, SemiBin2 requires only 11% of the peak memory usage of the SemiBin1 (see Supplementary Table 4). Therefore, applying SemiBin2 with multi-sample binning to large-scale metagenomic analysis will be much more efficient.

For long-read datasets, SemiBin2 also reconstructed the most near-complete bins (see Materials and methods). In the human gut, sheep gut and activated sludge projects with samples from different long-read sequencing technologies, SemiBin2 generated 13.2%, 28.1% and 14.8% more near-complete bins compared to the second-best binner (see Fig. 6). We also benchmarked these binners by evaluating the high-quality bins with 23S, 16S, 5S rRNA genes and tRNAs (see Materials and methods). When considering high-quality bins, SemiBin2 still performed the best and could reconstruct 15.6%, 26.3% and 13.1% more high-quality bins. Overall, SemiBin2 could outperform other binners in all situations for long-read datasets.

**Fig 6.**
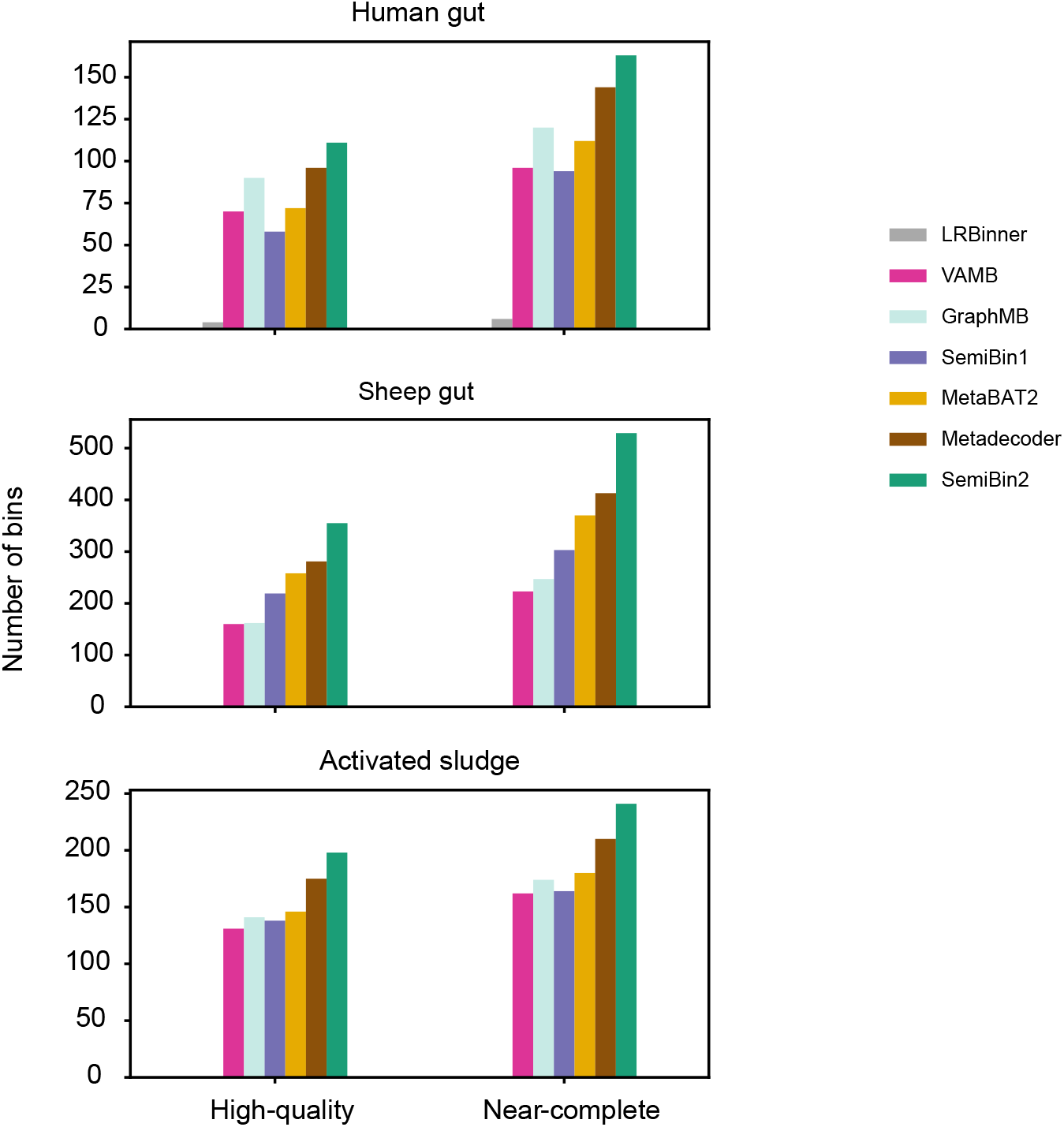
SemiBin2 outperformed other binners in long-read sequencing real datasets. SemiBin2 reconstructed more high-quality and near-complete genomes than other binners in three long-read sequencing projects. Near-complete bins: completeness > 90% and contamination < 5%; high-quality bins: completeness > 90%, contamination < 5% and having at least one 23S, 16S, 5S rRNA genes and 18 distinct tRNAs.

## 4 Discussion and conclusions

During the development of SemiBin1^20^, we found that contig annotations with GTDB taxonomy using MMSeqs2 performed better than annotations with the NCBI taxonomy using CAT^49^. Initially, our expectation was that continued improvement to reference genome databases and taxonomic prediction tools would lead to better binning results. However, there will always be novel species or strains in the environment^7^, and taxonomic prediction remains challenging and requires long running times. Thus, we attempted using random sampling to generate cannot-link constraints. Although there will be more noise in these randomly-sampled links compared to those from contig annotation (see Supplementary Table 2), deep learning can be robust to noise^47^. Empirically, cannot-link constraints generated by random sampling cover more genomes in the environment (including novel strains that the annotation algorithms cannot accurately identify), which leads to SemiBin2 getting better results than SemiBin1.

A similar idea to generate negative inputs has been used in CoCoNet^50^ for viral metagenome binning, which showed that the proportion of mislabeled contigs from the same genome is negligible. CoCoNet splits the contigs into fragments of size 1024bp to generate the cannot-link constraints, which is too short for metagenomic bacterial binning (contigs with this length are removed by most binning tools). SemiBin2 uses the whole contig for cannot-link constraints to learn a better embedding for clustering.

In recent long-read sequencing studies, MetaBAT2^16^, MaxBin2^17^, VAMB^19^ have been used^12^. However, these tools are optimized for short-read datasets. For this question, SemiBin2 uses an ensemble-based DBSCAN clustering algorithm for long-read datasets. SemiBin1 performed poorly in long-read datasets, as its community detection algorithm was not designed for long-read sequencing data.

This study proposes SemiBin2, a metagenomic binning method based on self-supervised learning. SemiBin2 outperforms other binners in both short- and long-read datasets (simulated and real). Compared to SemiBin1, by taking advantage of self-supervised learning, SemiBin2 requires much less running time and peak memory usage. Looking forward, there are other sources of information, such as the assembly graph (as used by GraphMB) and information from metaHiC related to physically linked contigs that can be considered to further improve binning results.

In addition to the algorithmic improvements discussed above, since the release of SemiBin1, we have also improved the tool in other ways. For example, SemiBin2 supports CRAM^51^ files, has more options for ORF finding, produces more statistics on its outputs, enables the user to better control output formats (*e.g*., filenames, and compression options) and—at the request of users—added support for re-using contig abundance estimates from MetaBAT2. In order to make it easier to run the tool as part of a pipeline, modules for nf-core^52^, Galaxy^53^ (available at the European Galaxy server, UseGalaxy.eu) and NGLess^54^ are now available. We have also fixed issues reported by users and improved error handling and reporting. Overall, SemiBin2 is more robust, easier to use and more flexible than SemiBin1, while returning better results at lower computational cost.

## 5 Acknowledgements

We thank Senying Lai (Fudan University) for her helpful comments on a previous version of this manuscript. We thank Bérénice Batut (University of Freiburg) for designing and implemementing the Galaxy wrapper. Members of the Zhao and Coelho groups are thanked for their comments throughout the development of this work. Users of SemiBin are thanked for their suggestions and bug reports.

## 8 Supplementary Figures and Tables

**Supplementary Fig 1.**
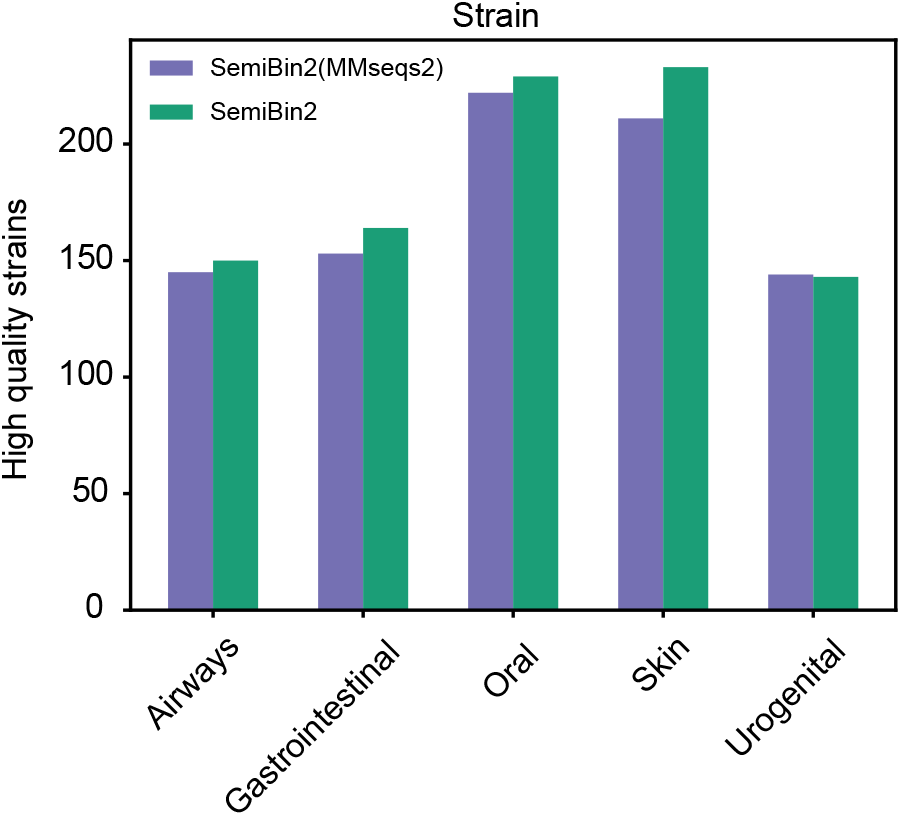
Self-supervised learning achieved better binning results compared to semi-supervised in CAMI II long-reads sequencing datasets. To compare the semi-supervised to self-supervised learning, we used their embeddings with the same ensemble-based DBSCAN clustering method in CAMI II long-reads sequencing datasets. Self-supervised learning achieved similar or better binning results compared to semi-supervised learning. SemiBin2(MMseqs2): using embeddings from semi-supervised learning; SemiBin2: using embeddings from self-supervised learning. Shown is the number of distinct high-quality strains.

**Supplementary Fig 2.**
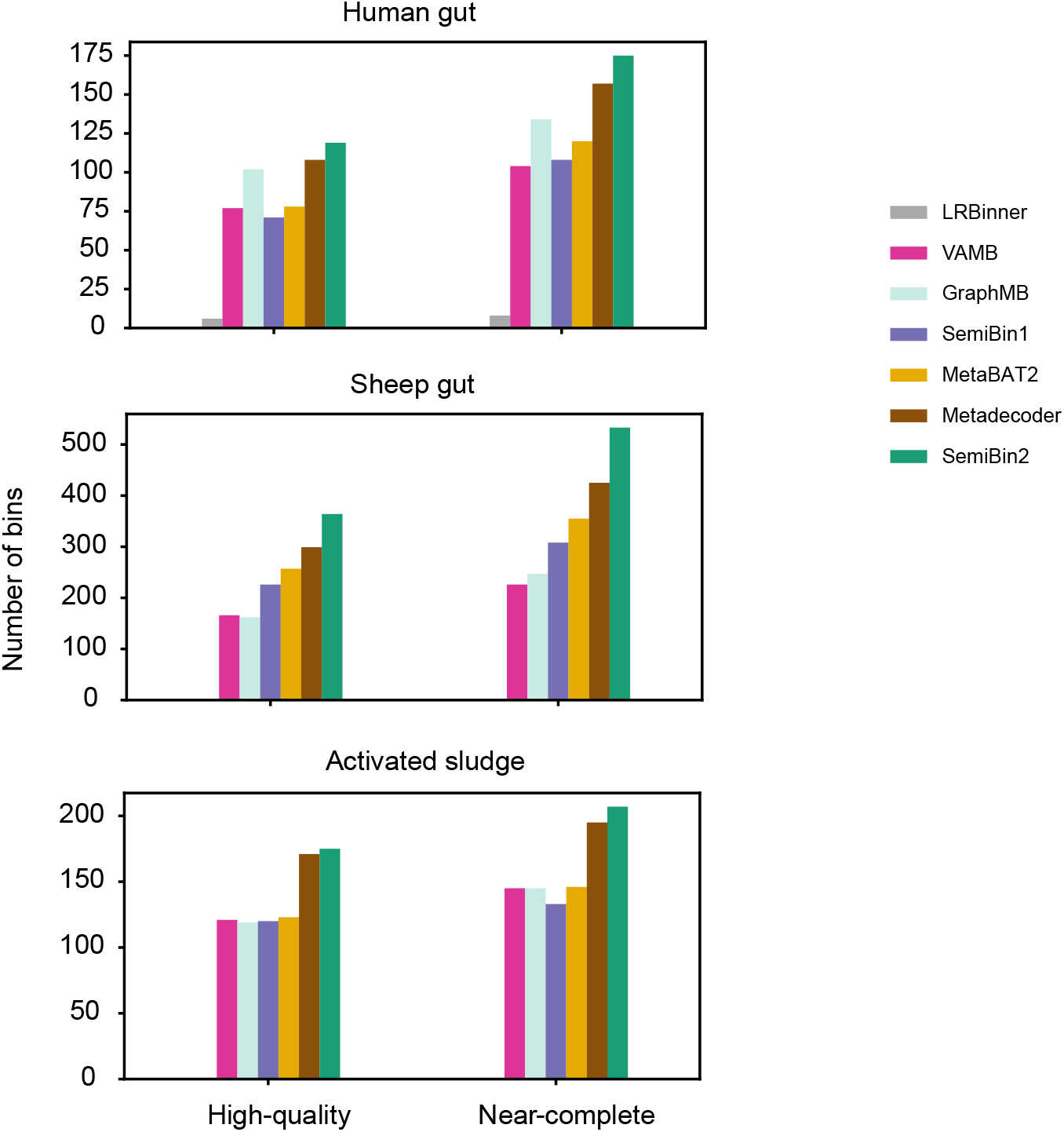
SemiBin2 outperformed other binners on real long-read datasets when performance was evaluated with CheckM1. As noted in the main text, because of the use of an overlapping set of genes for generating the bins and evaluation, this is not the strictest evaluation model.

**Supplementary Fig 3.**
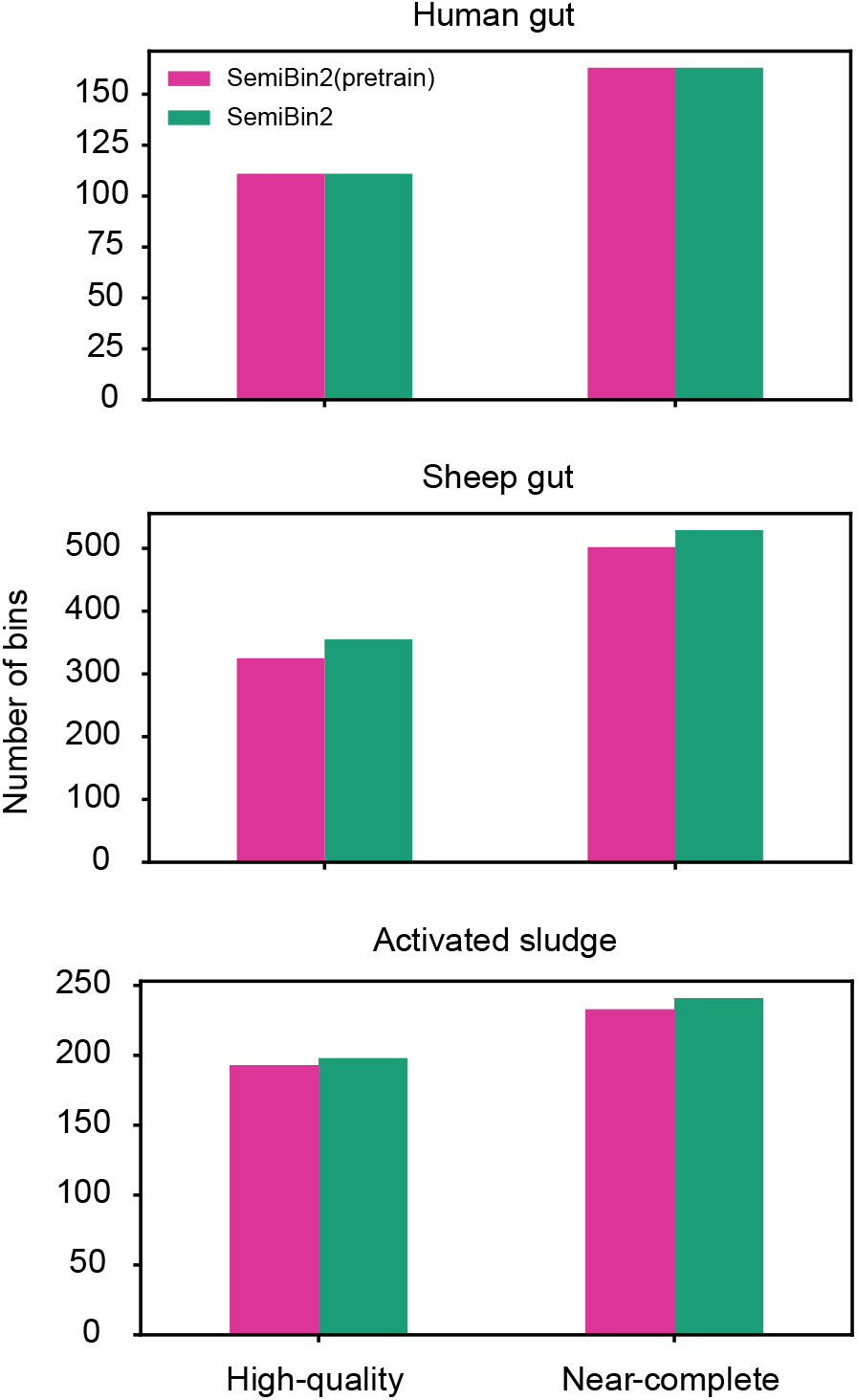
SemiBin2(pretrain) could also get good binning results on real long-read datasets. To test the performance of pretrained models on long-read datasets, we compared SemiBin2 (model trained from every sample) to SemiBin2(pretrain), which used the pretrained models already developed for SemiBin1^20^. For human gut dataset, we used the human_gut pretrain model and for other two projects, we used global pretrain model. Note that these pretrained models were built from short-read samples, they still produced good binning results on long-read datasets.

**Supplementary Table 1.** Statistics of assemblies from mNGS, PacBio-HiFi and Nanopore sequencing of 3 human gut samples. #contig: Number of contigs. Average length: average length of contigs. N50: the length of the shortest contig at 50% of the total assembly length.

|  |  | #contig | Average length | N50 |
| --- | --- | --- | --- | --- |
| Sample A | mNGS | 277,884 | 1,865 | 8,778 |
|  | PacBio-HiFi | 2,575 | 99,842 | 269,406 |
|  | Nanopore | 3,414 | 65,685 | 199,639 |
| Sample B | mNGS | 146,857 | 2,402 | 28,310 |
|  | PacBio-HiFi | 1,447 | 157,228 | 1,081,788 |
|  | Nanopore | 1,795 | 123,512 | 658,841 |
| Sample C | mNGS | 170,813 | 1,956 | 18,485 |
|  | PacBio-HiFi | 822 | 171,215 | 1,270,126 |
|  | Nanopore | 1,370 | 126,533 | 891,411 |

**Supplementary Table 2.**
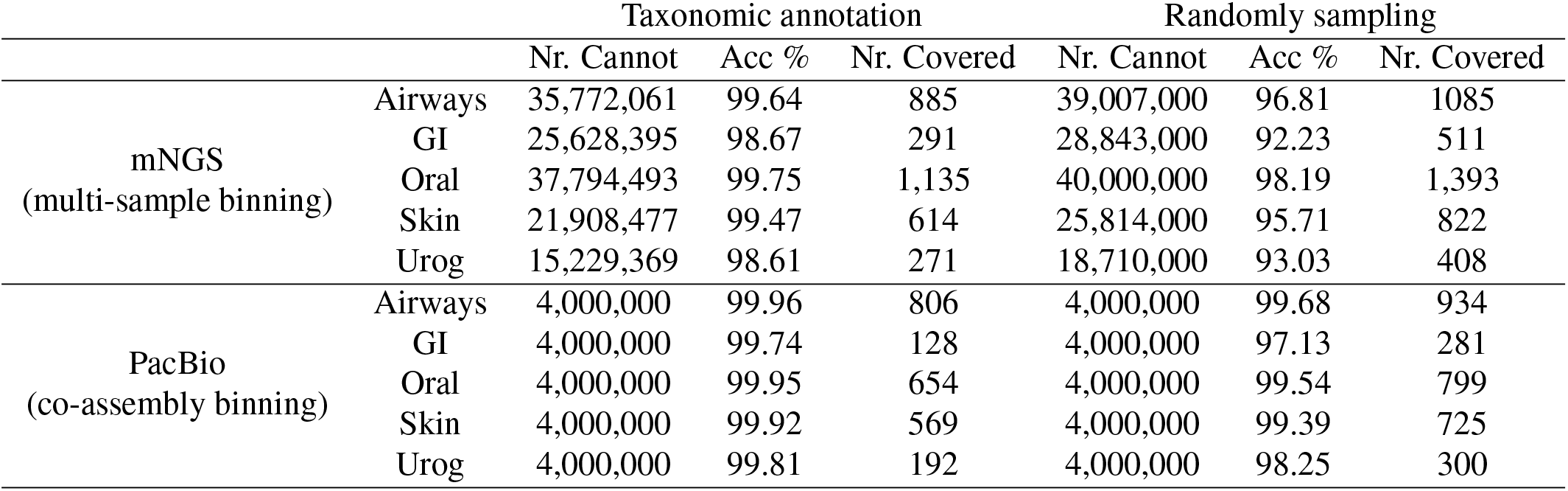
Accuracy of cannot-link constraints generated using MMseqs2 and randomly sampling in CAMI II simulated datasets. GI: Gastrointestinal. Urog: Urogenital. Nr. Cannot(MMseqs2): number of cannot-link constraints generated from MMseqs2. Acc(MMseqs2): the accuracy of cannot-link constraints generated from MMseqs2. Nr. Covered(MMseqs2): the number of genomes that are covered by the accurate cannot-link constraints generated from MMseqs2. Nr. Cannot(random), Acc(random) and Nr. Covered(random) are the corresponding results generated by randomly sampling.

|  |  | Taxonomic annotation |  |  | Randomly sampling |  |  |
| --- | --- | --- | --- | --- | --- | --- | --- |
|  |  | Nr. Cannot | Acc % | Nr. Covered | Nr. Cannot | Acc % | Nr. Covered |
| mNGS<br>(multi-sample binning) | Airways | 35,772,061 | 99.64 | 885 | 39,007,000 | 96.81 | 1085 |
|  | GI | 25,628,395 | 98.67 | 291 | 28,843,000 | 92.23 | 511 |
|  | Oral | 37,794,493 | 99.75 | 1,135 | 40,000,000 | 98.19 | 1,393 |
|  | Skin | 21,908,477 | 99.47 | 614 | 25,814,000 | 95.71 | 822 |
|  | Urog | 15,229,369 | 98.61 | 271 | 18,710,000 | 93.03 | 408 |
| PacBio<br>(co-assembly binning) | Airways | 4,000,000 | 99.96 | 806 | 4,000,000 | 99.68 | 934 |
|  | GI | 4,000,000 | 99.74 | 128 | 4,000,000 | 97.13 | 281 |
|  | Oral | 4,000,000 | 99.95 | 654 | 4,000,000 | 99.54 | 799 |
|  | Skin | 4,000,000 | 99.92 | 569 | 4,000,000 | 99.39 | 725 |
|  | Urog | 4,000,000 | 99.81 | 192 | 4,000,000 | 98.25 | 300 |

**Supplementary Table 3.** Overview of the datasets used in the benchmarking. GI: Gastrointestinal. Urog: Urogenital.

|  | Datasets | #Samples | #Genomes | Sequencing | Binning mode |
| --- | --- | --- | --- | --- | --- |
| CAMI II | Airways | 10 | 828 | mNGS | multi-sample |
|  | GI | 10 | 268 | mNGS | multi-sample |
|  | Oral | 10 | 799 | mNGS | multi-sample |
|  | Skin | 10 | 610 | mNGS | multi-sample |
|  | Urog | 9 | 254 | mNGS | multi-sample |
| CAMI II | Airways | 10 | 935 | PacBio | co-assembly |
|  | GI | 10 | 281 | PacBio | co-assembly |
|  | Oral | 10 | 799 | PacBio | co-assembly |
|  | Skin | 10 | 725 | PacBio | co-assembly |
|  | Urog | 9 | 301 | PacBio | co-assembly |
| Real datasets | Human gut | 82 | Unknown | mNGS | multi-sample |
|  | Dog gut | 129 | Unknown | mNGS | multi-sample |
|  | Ocean | 109 | Unknown | mNGS | multi-sample |
|  | Soil | 101 | Unknown | mNGS | multi-sample |
| PRJCA007414 | SAMC515478 | 1 | UnKnown | PacBio-HiFi | single-sample |
|  | SAMC515479 | 1 | UnKnown | PacBio-HiFi | single-sample |
|  | SAMC515480 | 1 | UnKnown | PacBio-HiFi | single-sample |
|  | SAMC515478 | 1 | UnKnown | Nanopore R9.4 | single-sample |
|  | SAMC515479 | 1 | UnKnown | Nanopore R9.4 | single-sample |
|  | SAMC515480 | 1 | UnKnown | Nanopore R9.4 | single-sample |
| PRJNA595610 | SRR10963010 | 1 | UnKnown | PacBio-HiFi | single-sample |
|  | SRR14289618 | 1 | UnKnown | PacBio-HiFi | single-sample |
| PRJEB48021 | SAMEA9994818 | 1 | UnKnown | PacBio-HiFi | single-sample |
|  | SAMEA9994819 | 1 | UnKnown | NanoPore R9.4.1 | single-sample |
|  | SAMEA9994820 | 1 | UnKnown | NanoPore R9.4.1 | single-sample |
|  | SAMEA9994714 | 1 | UnKnown | NanoPore R10.4 | single-sample |

**Supplementary Table 4.**
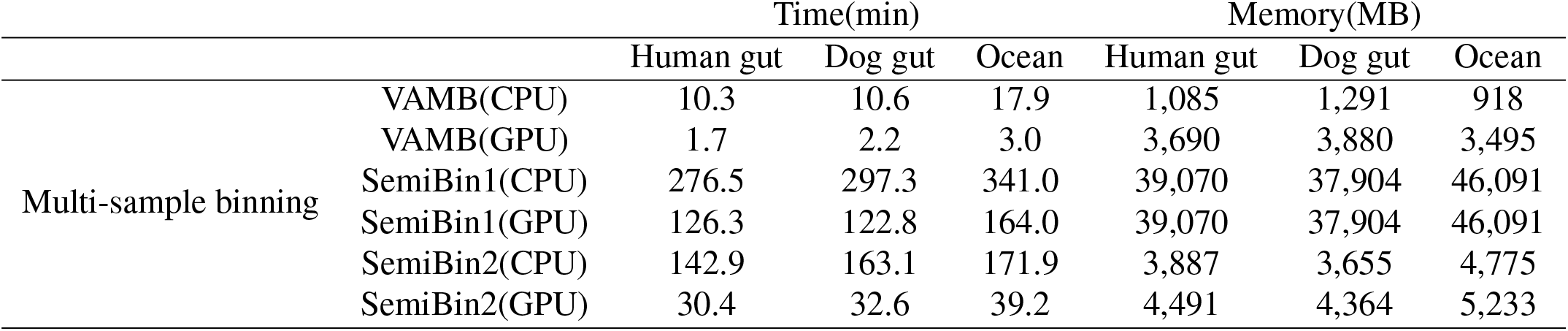
Running time and memory usage for the different binning tools with multi-sample binning. Average per sample time and peak memory usage (over 10 randomly chosen samples). For CPU timings, we used an AWS g4ad.4xlarge machine with 1 CPU, 8 physical cores and 16 logical cores. For GPU machine, we used a Tesla T4. GPU: Graphical Processing Unit, CPU: Central Processing Unit.

|  |  | Time(min) |  |  | Memory(MB) |  |  |
| --- | --- | --- | --- | --- | --- | --- | --- |
|  |  | Human gut | Dog gut | Ocean | Human gut | Dog gut | Ocean |
| Multi-sample binning | VAMB(CPU) | 10.3 | 10.6 | 17.9 | 1,085 | 1,291 | 918 |
|  | VAMB(GPU) | 1.7 | 2.2 | 3.0 | 3,690 | 3,880 | 3,495 |
|  | SemiBin1(CPU) | 276.5 | 297.3 | 341.0 | 39,070 | 37,904 | 46,091 |
|  | SemiBin1(GPU) | 126.3 | 122.8 | 164.0 | 39,070 | 37,904 | 46,091 |
|  | SemiBin2(CPU) | 142.9 | 163.1 | 171.9 | 3,887 | 3,655 | 4,775 |
|  | SemiBin2(GPU) | 30.4 | 32.6 | 39.2 | 4,491 | 4,364 | 5,233 |

